# A computational workflow for microscopy-guided ion identification in clinical mass spectrometry imaging datasets

**DOI:** 10.64898/2025.12.25.694368

**Authors:** Ana-Maria Năstase, Ping K. Yip, Christopher E.G. Uff, Irma O’Meara, Hervé Barjat

## Abstract

Matrix-assisted laser desorption/ionisation mass spectrometry imaging (MALDI-MSI) datasets were acquired from brain tissue samples obtained from living traumatic brain injury (TBI) patients. This is a proof-of-concept study which presents a computational workflow for identifying TBI-specific ions in MSI datasets through integrated pathological annotation and mass spectrometry imaging analysis. Pathological annotations of TBI regions were obtained from haematoxylin and eosin-stained microscopy images of the same sections. The microscopy images were then registered with the MSI datasets to enable precise delineation of TBI areas within the MSI data. Binary masks of the registered TBI regions were then used to perform cosine similarity searches, identifying ions potentially associated with TBI pathology. In order to relatively quantify the difference magnitude between TBI regions and surrounding tissue, bootstrap sampling was applied to non-TBI tissue areas, generating mean intensity values for comparison with average TBI region intensities. Additionally, a synthetic MSI dataset was generated to validate and optimize this analytical approach. Using this integrated computational workflow, TBI specific ions were identified in clinical MSI datasets.

## Introduction

Mass spectrometry imaging (MSI) is a technique in which mass spectra are first acquired at defined positions across the two dimensional surface of a sample and then reconstructed as ion distribution maps, also known as ion images [39], reflecting molecular distributions across the sample. MSI data analysis workflows can vary depending on the objective of each experiment. When it comes to studying complex biological systems, such as tissue from biopsies, the delineation of regions of interest (ROIs) is key to understanding the underlying biochemical layout within the sample. Spatial segmentation is one of the ROI selection strategies capable of identifying the molecular heterogeneity of a specimen. Spatial segmentation can be elucidated using dimensionality reduction techniques followed by clustering algorithms [15]. This technique is recommended for exploratory analysis. When interested in investigating the biology of specific tissue areas, however, ROIs delineation guided by histopathological information leads to a more accurate data analysis. This is particularly important for the application of MSI in clinical studies, where MSI data sets and histological annotations can be used for further pathological investigation through data analysis based on ROI delineation [12–14, 16, 36, 37, 50, 51].

In this study, pathological annotations of histopathology images were used to guide the ROI selection. Microscopy images of tissue cryosections stained with haematoxylin and eosin (H&E) were generated and annotated independent from the MSI data. This method ensures an accurate identification and delineation of ROI in the MSI dataset in a non-biased approach [13, 14, 16, 50]. Given that this multimodal approach was employed, co-registration between the two modalities was essential to ensure that H&E annotations accurately transfer to the MSI dataset. After the ROI has been delineated in the MSI dataset, the analysis focused on developing downstream ROI extraction process. This method aims to identify ions with ROI-specific signal, i.e. ions that are specifically present in a particular ROI. In this study, we propose using the ROI binary mask to aid in the identification of ions associated with the ROIs obtained from the pathological annotations. The aim was to develop a simple computational workflow which extracts the ions associated only with specific tissue regions from a clinical MSI dataset.

For determining the spatial pattern match between the ion images and the ROI binary masks, cosine similarity was used. This method is widely used in the MSI field for identifying co-localized ions, where one of the ions of interest is used as the reference image. A recent study analysed different similarity measures and algorithms for ion co-localization and determined that among the classical approaches the cosine similarity was a good similarity metric for comparing MSI images [30]. Cosine similarity is widely applied in multiple different fields, including topic modeling, anomaly detection, and image similarity because it can measure the angle between two vectors regardless of their magnitude [19]. This metric is a measurement of orientation and not of magnitude and it is useful for sparse data where zero values do not necessarily indicate a lack of similarity. In MSI, studies focus on ion-to-ion comparison with cosine similarity, rather than ion-to-binary mask [18, 27, 30]. Therefore, this approach proposes an extension of existing methods, as it adds the capability to search within high dimensional MSI data for ions matching specific spatial patterns of interest.

There are several advantages of using cosine similarity scores with the ROI binary mask. Firstly, cosine similarity handles sparse data well. Secondly, as it measures the angle between the vectors, it is suitable for use in pattern similarity matching. Finally, it is unaffected by the differences in vector magnitudes. Moreover, due to the lack of an objective control region within the sample and inherent heterogeneity of the sample, a pattern search is advantageous.

Since the spatial pattern similarity search yields no information regarding average intensity differences between the different regions within the tissue, possibly due to masking of selective micro-regions with a gross background activity, it needs to be complemented by a method that measures average region intensity differences between ROI and control regions [44]. This was achieved by comparing the mean intensities of the pixels in the two regions from the biopsy samples: uninjured control, i.e. tissue area excluding ROI, and ROI. Due to the fact that there were no specific control regions within the tissue in the present work, and considering that the control area is generally much larger than the ROI area, random pixel samples from the control area equally sized to the ROI area were created. This sampling process is similar to the one applied in bootstrap sampling, a method that makes no assumptions regarding the underlying data distribution [10]. This “hybrid” bootstrap sampling approach focuses on the ROI as the investigation target, while also controlling for the sample size bias. Another advantage of using this method is that it provides uncertainty estimates which is particularly useful for MSI data where spatial heterogeneity is frequently encountered [37].

This workflow, consisting of image registration, cosine similarity search and “hybrid” bootstrap sampling approach, was successfully employed in the analysis of a dataset acquired from brain tissue samples obtained from living traumatic brain injury (TBI) patients. TBI refers to the brain damage caused by external forces such as falls from height, road traffic collisions, and assaults [4]. Different mechanisms are involved in the pathophysiological response to TBI such as synaptic dysfunction, mitochondrial dysfunction and neuroinflammation [42]. There is currently no FDA approved therapy for moderate or severe TBI [42]. Therefore, there is a need to improve diagnosis and prognosis through the elucidation of TBI mechanisms in the various forms of severity and, thus, for biomarker discovery [8, 11, 26, 42]. A few recent studies have employed mass spectrometry (MS) techniques to determine protein biomarkers in biofluids of living and post-mortem patients with TBI [5, 7, 40, 52]. However, molecular imaging studies have mostly focused on assessing translational biomarkers of TBI in animal models [1, 20–22, 24, 25, 29, 41, 43] and no studies have looked to focus specifically on the spatial peptide biomarkers found only in the TBI regions of brain tissue biopsies from living TBI patients. More specifically the workflow was aimed at obtaining a list of possible peptides that could be related to the biology of TBI.

## Methods

### Workflow

The full workflow of the study (Figure 1) was comprised of multiple parts including MSI data processing, H&E images annotation, co-registration between the two modalities followed by ROI cosine similarity search and difference calculation between disease and control regions using bootstrap sampling [9], each of them being fully detailed in this section. The clinical TBI dataset was processed using Fiji [34, 35] (ImageJ, NIH) and Cardinal [2, 3] (running on R [32]) and in-house developed Python-based tools (version 3.11.13, [46]). The terms ion and mass channel are used interchangeably throughout.

**Figure 1.**
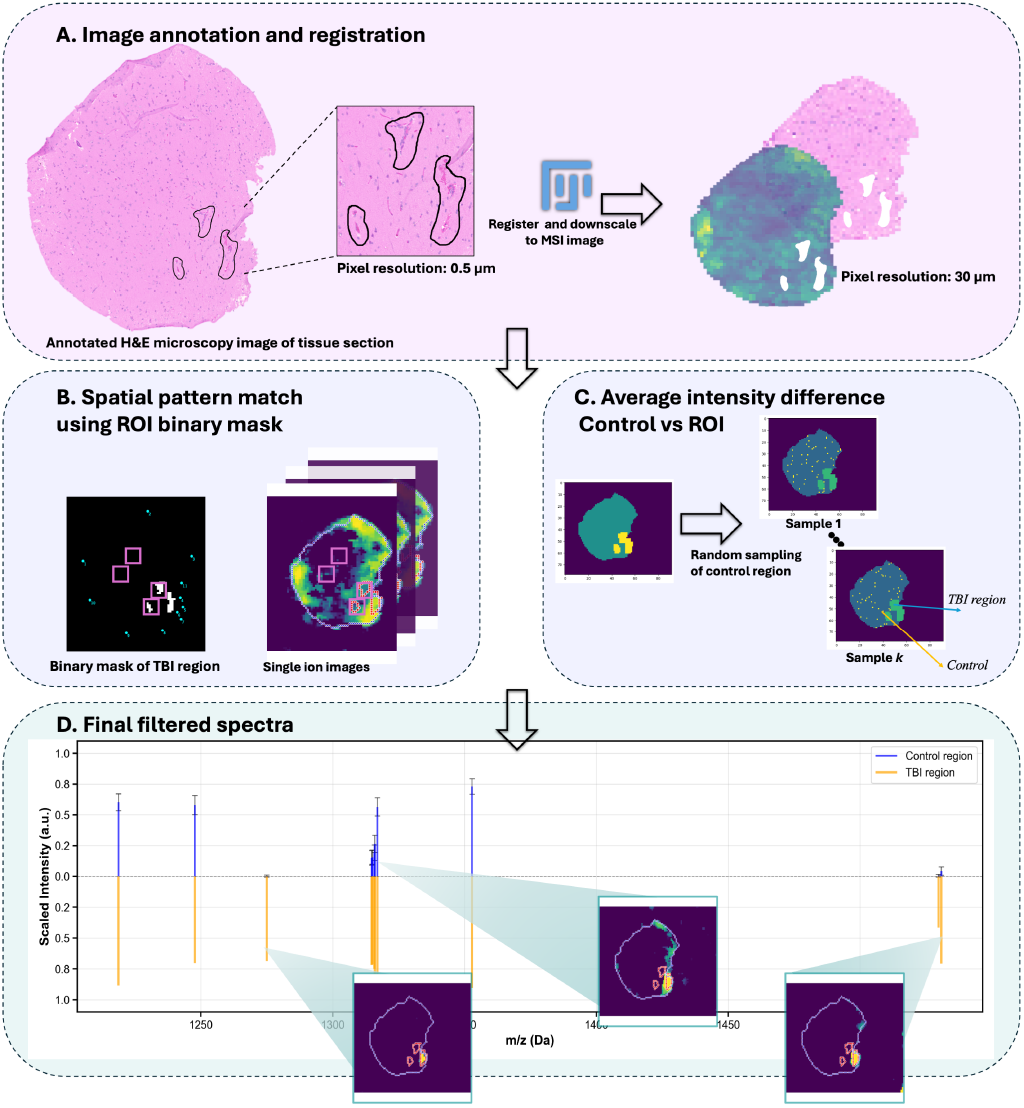
Diagram of the workflow developed for detecting ROI-specific mass channels in MALDI-MSI data. A. Image annotation and registration: This part focused on the image registration between microscopy and MSI images using Fiji. At this stage the microscopy images were also annotated for ROIs blind to the MSI data. B. Spatial pattern match using the ROI binary mask. The binary mask for the ROI was used to calculate the cosine similarity score for each mass channel in the TBI dataset. C. Average intensity difference. The control region was randomly sampled multiple times to obtain an average and compared with the TBI average. D. Final filtered spectra. The top most similar ions to the binary mask were selected and plotted.

### Data pre-processing

In total, 24 clinical brain tissue biopsy samples were analysed and used to develop the workflow presented in this manuscript. The biopsy samples were collected from living patients who suffered a TBI as part of the Severe Head Injury Brain Analysis (SHIBA) study, IRAS ID 271662 [45] and were provided by Queen Mary University of London (QMUL) and Barts Health NHS Trust. The dataset itself is confidential and only certain technical aspects regarding it can be disclosed – *m/z* values and annotations are not presented, as this is outside the scope of this manuscript. Detailed information on the sample preparation and acquisition using matrix-assisted laser desorption/ionisation (MALDI)-MSI platform at 30.3 *µm* may be found in the Supporting Information (SI 1.1).

The raw imzML files [33] were obtained from the flexImaging software (Bruker, Germany). The MALDI datasets were all pre-processed using Cardinal (R library) by centroiding and data normalisation. Data centroiding which also incorporates baseline removal, Gaussian smoothing, and pixel mass-to-charge ratio (*m/z*) alignment was performed. The rest of the analysis was carried out in Python where the data was first normalised using median calculation followed by logarithmic (base 2) transformation.

### H&E microscopy and annotations

Following the MALDI-MSI analysis, the tissue sections were subsequently stained with haematoxylin and eosin (H&E) and images were acquired using a microscopy platform at 0.55 *µm* (SI 1.4). The microscopy images were annotated using Fiji using the ROI manager. TBI ROIs, often indicated by micro-haemorrhage, were defined manually by a central nervous system (CNS) neuroinjury expert using the freehand selection tool, without using information from the corresponding MSI data. Multiple ROIs could be present in one sample. A binary mask of the ROI regions was generated in Fiji for each sample (Figure 2). Functionality was wrapped up into macros.

**Figure 2.**
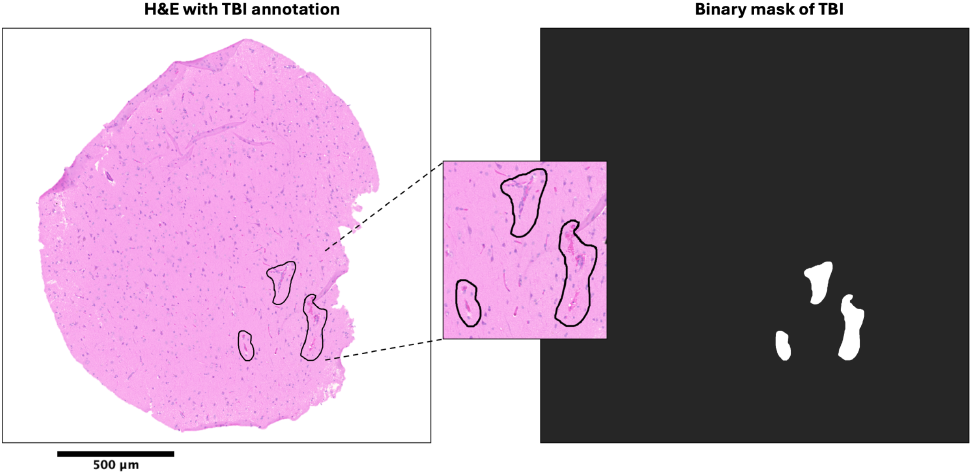
Example of H&E microscopy (left) of brain tissue section with delineated TBI regions alongside its binary mask (right).

### MALDI-MSI and microscopy co-registration

Fiducial markers were used to unambiguously determine the image orientation across the two imaging modalities (SI Figure 8). The co-registration between MALDI-MSI and microscopy was also performed semi-automatically in Fiji. Given the difference in image resolution between the two methods, landmark correspondences (TransformJ [28]) was used to ensure accurate elastic registration between the two images. This process is similar to the one described in [31]. Since the aim was to map the ROI binary mask from the microscopy image on to the MSI image, the former image was set as the moving image (source image) and the latter image as the fixed one (template image). The transformation method used was moving least squares (non-linear) and the transformation class affine.

In order to select the landmarks which best delineate the tissue in the MSI dataset, for each sample a MSI image stack containing single ion images, segmented images and total ion count images was generated. Due to the low ion signal intensity coming from the tissue, a combination of these images was necessary in order to be able to accurately pinpoint the tissue landmarks used for the elastic image registration. Varying numbers of landmarks were selected for each sample, depending on the size and quality of the data.

For this study three areas per tissue were selected, the TBI region (TBI), the control region which was the whole tissue region without the TBI (WT) and the background region (BG). The binary masks for the regions of interest from the annotated microscopy data were also transformed using the same transformation matrix.

### Cosine similarity search using the ROI binary mask

Let us consider our MSI dataset *M* (∈ ℝ^*m*×*n*^), where *m* is the total number of mass channels and *n* the total number of pixels and *M*_*ij*_ refers to mass channel *i* at pixel/spectrum *j*. Each *M*_*i*_ was flattened to a 1D vector and scaled to [0,1].

For the downstream analysis, to ensure the signal is coming only from the tissue and not from the background, as part of this step, for *M*_*i*_ a mean BG spectrum intensity 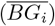 was also calculated and subtracted from the mean WT spectrum intensity 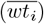 If for *M*_*i*_ the 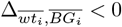 then, *M*_*i*_ is filtered out.

This method focused on image co-localisation using only the binary TBI mask notated as *B* below. This ensures that the ions are spatially co-localised with the disease region. This method was performed using cosine similarity for calculating the distance metric between each mass channel image and the TBI mask. This is defined by the following formula:

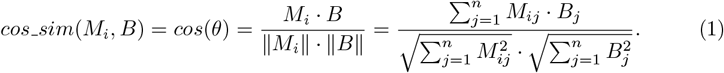

The similarity score for all mass channels is stored as a vector *Cos_sim* = [*cos_sim*(*M*_1_, *B*), *cos_sim*(*M*_2_, *B*), …, *cos_sim*(*M*_*m*_, *B*)] ∈ ℝ^*m*^. In practice, SciPy package [47] was used to determine the cosine distance which was then converted to cosine similarity as follows: *cos_sim*(*M*_*i*_, *B*) = 1 – *cos_dist*(*M*_*i*_, *B*).

### Average ROI intensity difference using bootstrap sampling

The next method focused on the average pixel intensity comparison between the TBI and WT regions for each mass channel in an MSI dataset. This method uses random sampling from the WT region to obtain the WT average and compare it to the TBI average for each resampling step. This method recognises the asymmetric nature of the two regions in terms of sample size, and also provides uncertainty information for the comparison of interest. This workflow was developed in Python.

This method follows a similar process to bootstrap sampling. Consider two subsets of dataset *M*, one for the control region *C*(∈ ℝ^*m*×*q*^) and one for the TBI region *T* (∈ ℝ^*m*×*r*^), where *q* is the total number of pixels in the control area and *r* is the total number of pixels in the TBI area. The control area refers to the WT area minus the dilated TBI region. A new region corresponding to the dilated TBI region was created and excluded from the control area as the biology in the periphery of the TBI region is likely to be quite variable from region to region and sample to sample. This is also known as a penumbra and is generally observed in TBI [48]. This was performed using OpenCV [6] by applying morphological transformations with a rectangular structuring element. Considering the number of pixels added after the dilation *d* and the total number in the WT area *l*, then the number of pixels *q* in the “safe control region” is defined by *q* = *l* − (*r* + *d*).

The random sampling is performed from *C* and the bootstrap sample for each mass channel is *C*_*ik*_ = [*x*_1_, *x*_2_, …, *x*_*r*_] and *K* = [1, 2, …, *k*] is the number of iterations for the resampling process. For each iteration *k*, for any *M*_*i*_ the difference between 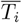 and 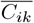 is calculated, which results in vector 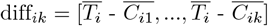. This allows for the subsequent determination of additional statistical measures such as mean, standard deviation and confidence interval of diff_*ik*_. For each sample, a *t*-test may be performed to determine whether there is any statistically significant difference between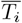 and 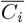.

### Generating a synthetic dataset

A synthetic dataset was generated using the masks of one of the samples in the TBI dataset. The control mass channels were generated using only the WT binary mask by assigning a random intensity value between [1,5] while adding or subtracting a random number of blobs of varying blob-to-tissue intensities (b:T). This was done in order to ensure the tissue signal replicates the heterogeneity generally encountered in clinical data. In the case of the synthetic dataset analysis the equivalent of the TBI region is notated simply as ROI. The ROI specific mass channels were generated similarly to the control mass channels with the addition of upregulated or downregulated ROIs with different ROI-to-tissue intensities (R:T). Finally, Poisson noise which is characteristic to MSI images was added to all mass channels [38]. The *m/z* values were randomly generated in the interval [700,1800]. These random *m/z* values will be referred to as channel IDs. The intensity of the blobs and ROI regions was Gaussian distributed with a pre-determined covariance matrix.

## Results and Discussion

### Image registration

The resolution of the H&E microscopy images is 0.55 *µm*, which means that the area of a pixel is 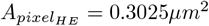 In contrast, the resolution of the MSI images in the clinical TBI dataset is 30.3 *µm*, i.e. the area of an MSI pixel is 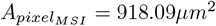 This roughly translates to 1 pixel in MSI being equal to around 3040 pixels in H&E. Hence, any annotated ROIs with an area smaller than 3000 pixels in the H&E image were not included in the analysis. Binary masks from registered H&E images were generated for each sample for both the TBI regions and the WT regions.

The results from one of the samples from the clinical TBI dataset are presented next to showcase the application of the workflow presented in Figure 1. Figure 4 highlights the landmark-based registration process performed in Fiji between the H&E microscopy image and one of the images in the MSI stack, which in this case was the image of the first principal component of the sample dataset. Both images have two overlays originating from the WT mask and the TBI mask. In the case of the MSI the overlays originating from the registered masks illustrate the quality of the image registration.

### Synthetic dataset - cosine similarity

The synthetic dataset was generated using the mask shown in Figure 3a, with examples of synthetic ion images being provided in SI Figure 9. Control mass channels were set at 50% of the total and ROI-specific mass channels were divided into five categories focused on ROI regulation patterns: three upregulated categories with varying ROI-to-tissue intensity ratios (R:T of 3, 5, and 10), and two downregulated categories with tissue-to-ROI ratios (T:R) of 3 and 10. The cosine similarity method (Figure 1B) was then evaluated on this dataset.

**Figure 3.**
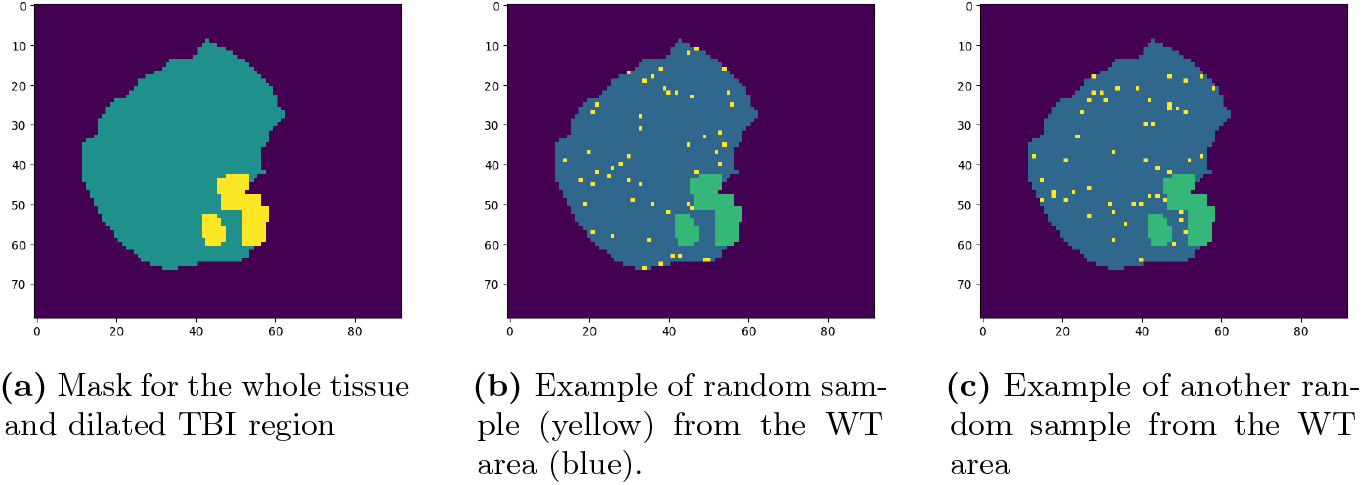
Figures showcasing the bootstrap sampling performed. 3a shows the mask for the TBI dataset (TBI area-yellow and WT - green). 3b and 3c show two examples of random samples (yellow) obtained from the WT area, where the size of the sample is equal to the TBI region size

**Figure 4.**
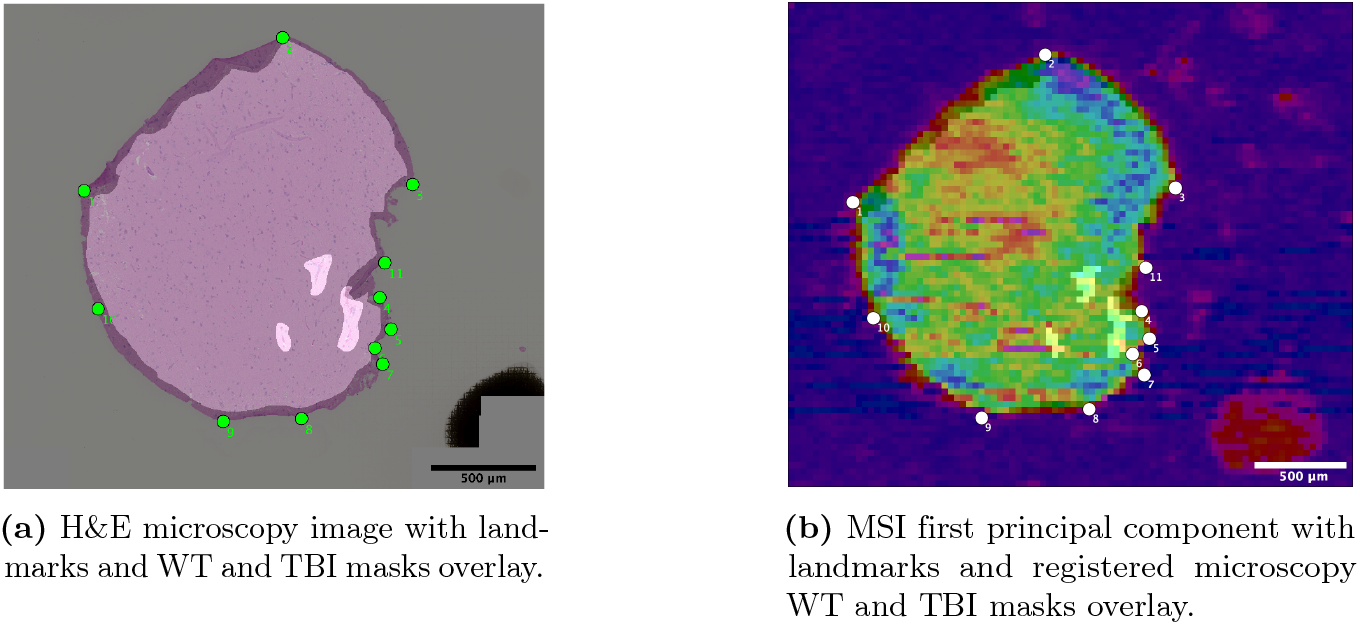
Registration results for sample 9 with landmarks shown as numbered points which illustrate the one-to-one correspondence between panels 4a and 4b. Note that in this dataset the fiducial marker used to determine image orientation is visible in the bottom right corner of the images.

Figure 5a displays the cosine similarity score assigned to each mass channel. Control mass channels were represented with blue dots and mass channels with upregulated or downregulated ROI were represented with red or green dots, respectively. The purple horizontal line indicates the average score of the control mass channels. Upregulated ROI mass channels generally have a higher similarity score than the control mass channels, while the downregulated ROI mass channels appear to have a lower similarity score than the control mass channels (Figure 5a). This trend is maintained even if an inverse similarity score (calculated by using the inverse of the scaled ion image) is used for downregulated ROI mass channels. Since these scores do not provide a clear threshold for selecting ROI-specific mass channels which also include downregulated ROI mass channels, absolute *z*-scores derived from these scores (Figure 5b) could be used to select a cut-off value which leads to the inclusion of downregulated ROI in the filtered data (Figure 1D). Measures of observational errors such as precision and accuracy were not included, as no objective cut-off value could be identified. However, if downregulated ROI mass channels are not of interest, or not biologically relevant, the *z*-scores would not be needed and only the similarity scores could be used. Figure 5 also points to an influence of the R:T on the similarity score, where mass channels with higher R:T have a higher cosine similarity score.

**Figure 5.**
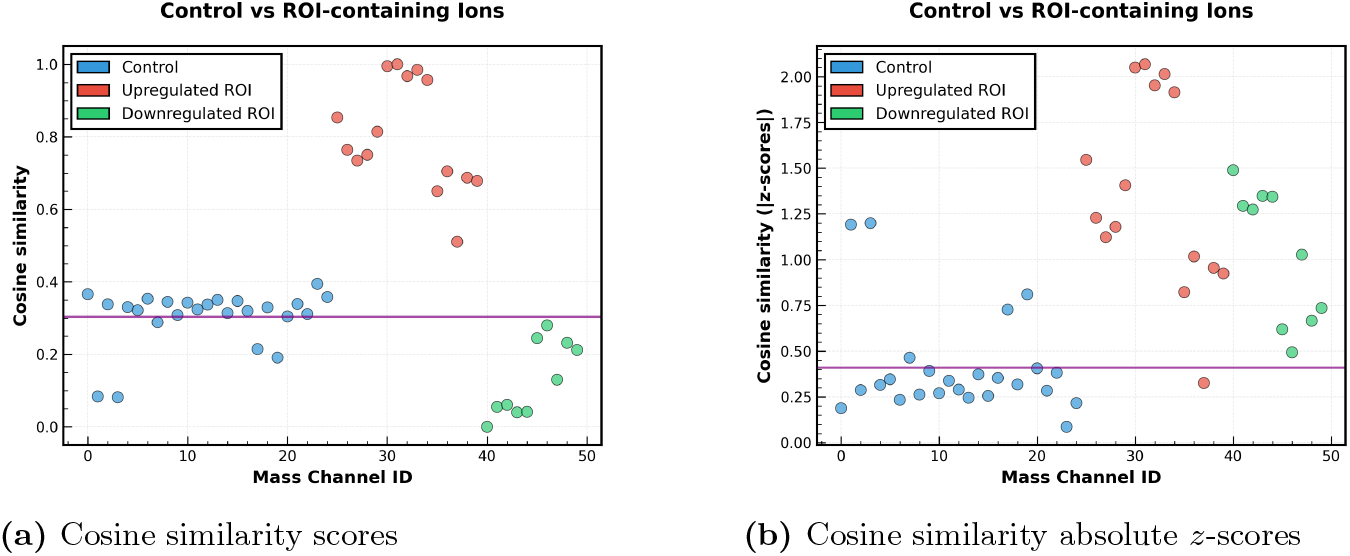
Cosine similarity scores for the synthetic MSI dataset. The dataset includes both control and ROI-specific mass channels. The ROI-specific mass channels were generated so that the average intensity was either upregulated or downregulated. The mean value of the control mass channels is marked with a horizontal purple line.

In conclusion, the cosine similarity method successfully identified upregulated ROI mass channels. However, downregulated ROI mass channels showed a lower chance of being identified as ROI-specific compared to control mass channels (Figure 5).

### Clinical TBI dataset results

The mirror plots in Figures 6 and 7 display the top 0.5 percentile mass channels based on *cos_sim z*-scores, while also highlighting the differences in the average intensity between the two regions (diff_*i*_). The percentile value could be modified to broaden the search of relevant *m/z* values. In this case, a stringent percentile value was selected for visualisation purposes.

**Figure 6.**
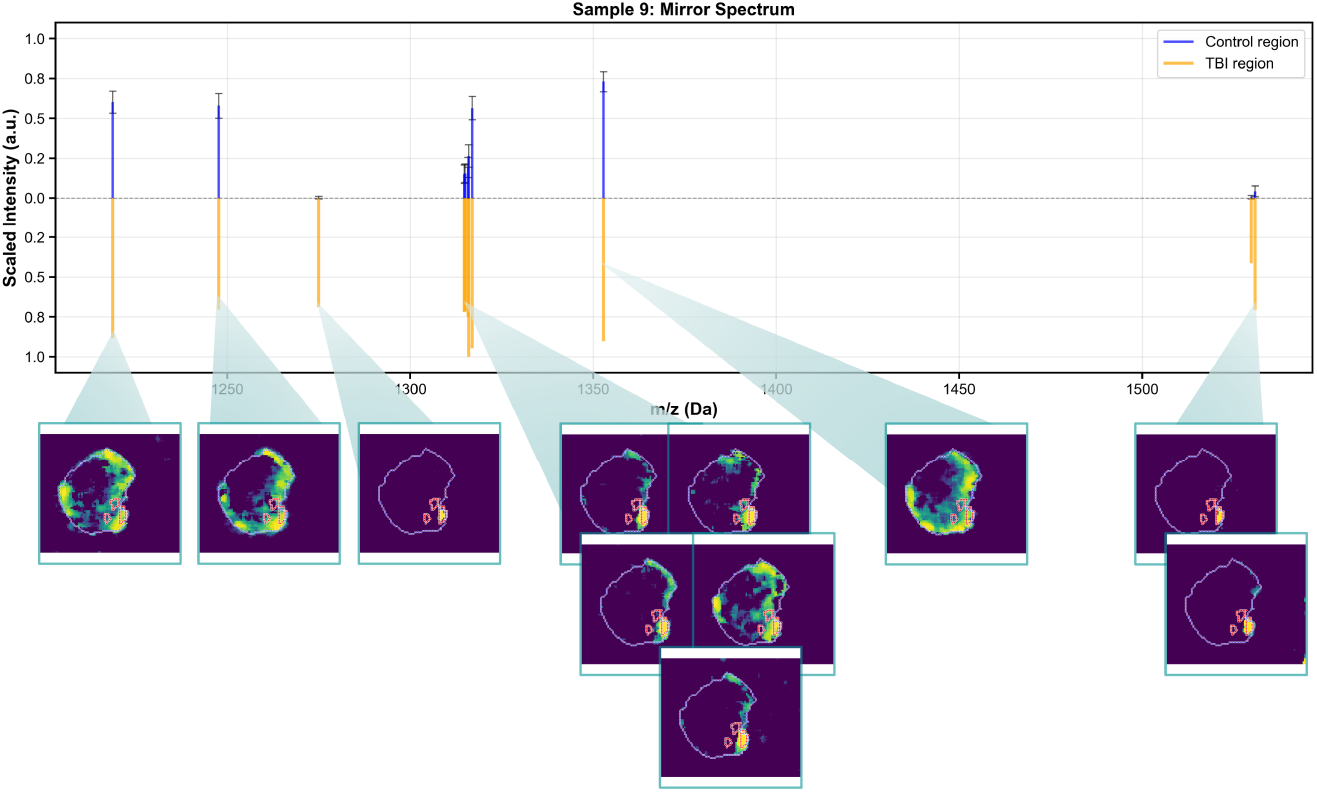
Mirror spectrum of the top percentile ions filtered based on their *cos*_*sim*(*M*_*i*_, *B*). The control spectrum *C*_*i*_ is presented with the standard deviation obtained from the bootstrap sampling. For each ion image, the WT area is marked with purple contour and the TBI region is marked with red contour.

**Figure 7.**
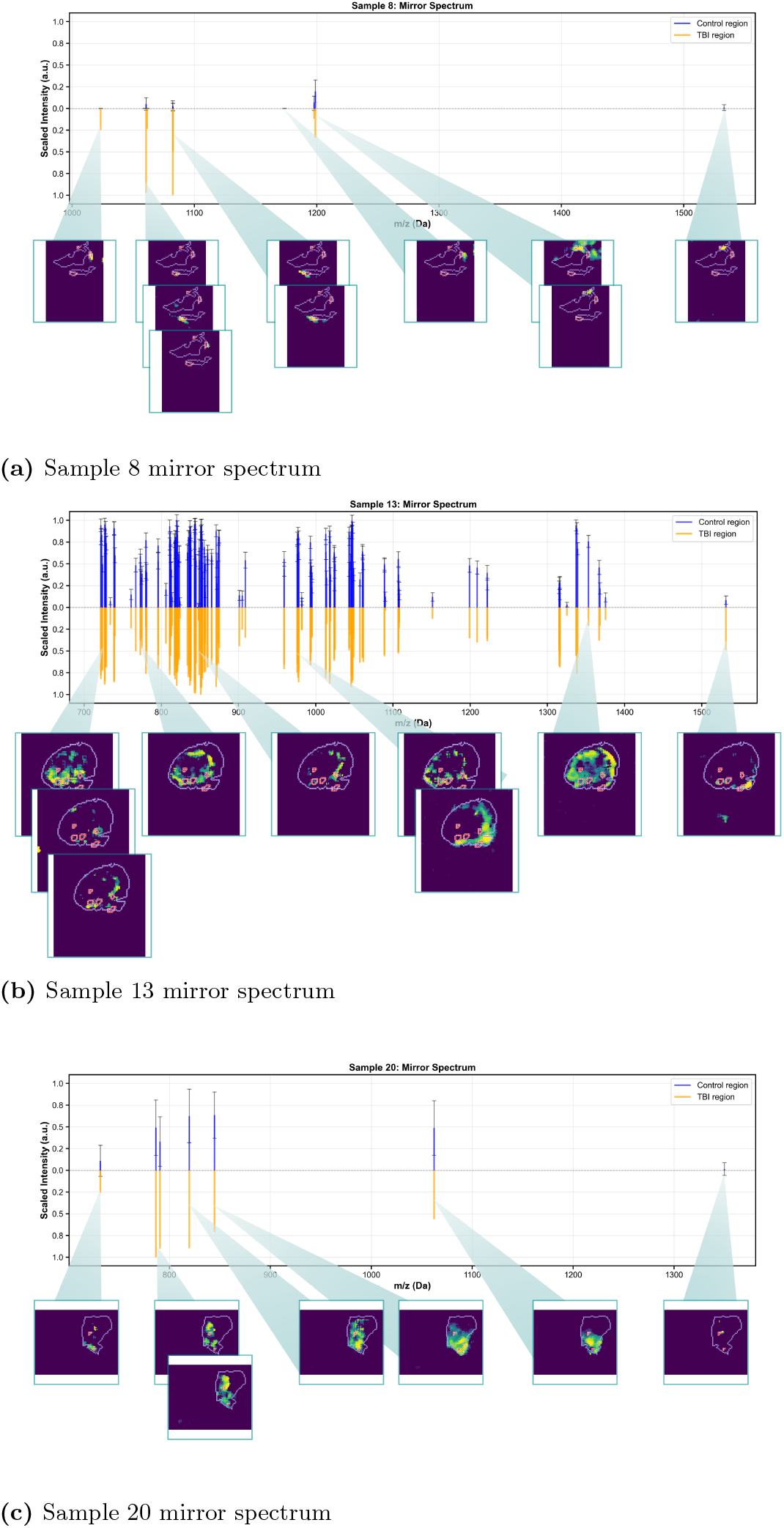
Further examples of mirror spectra from the TBI dataset.

**Figure 8.**
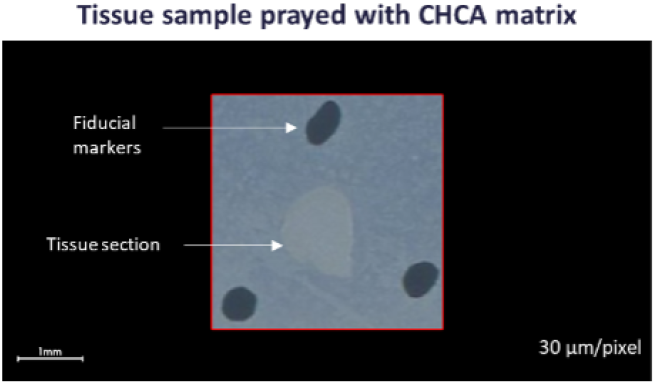
Example trypsin-digested TBI sample sprayed with CHCA matrix, showing the fiducial markers added around it.

**Figure 9.**
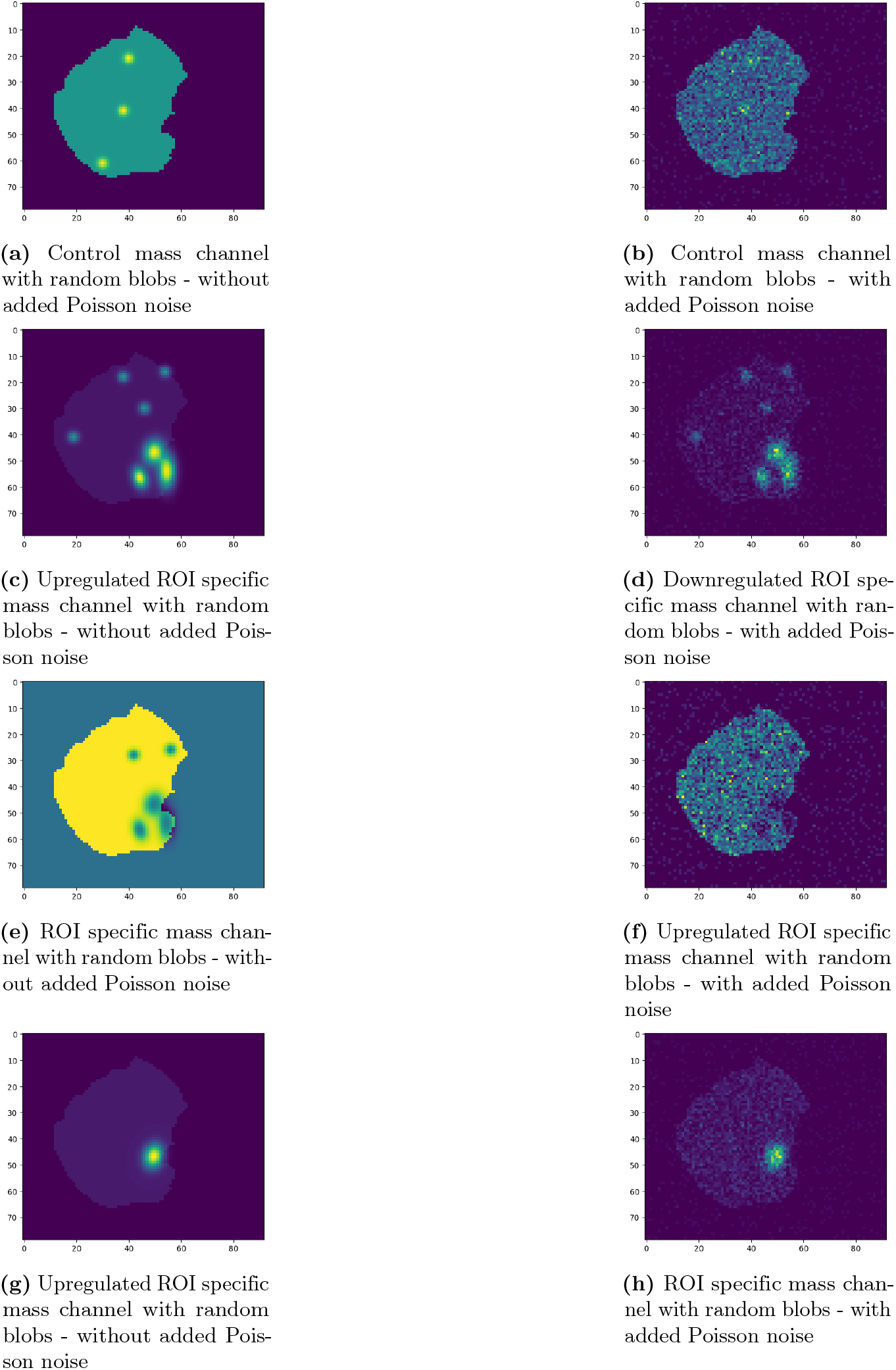
Examples of mass channels generated for the synthetic dataset.

Figure 6 shows results for the sample used to demonstrate the workflow application. The control average intensity (*C*_*i*_) was calculated using the bootstrap sampling process (Figure 1C). All filtered ions in this sample exhibited upregulation in the ROI (diff_*i*_ *>* 0). Moreover, for some of the ions, the signal coming from the TBI region appears to extend into adjacent areas (peri-injury areas).

Additional examples of filtered ions are shown in Figure 7 for the purpose of discussing the results obtained from this workflow. In Sample 8 (Figure 7a), one ion shows a diff_*i*_ and standard deviation of zero, yet it ranks in the top percentile based on its *cos_sim*(*M*_*i*_, *B*). The corresponding ion image below the spectrum reveals that the signal originates just outside the delineated TBI and WT regions. This discrepancy may arise from several factors such as registration errors or inaccurate ROI annotation. Registration images for this particular sample are supplied in SI Figure 10. If the signal is indeed coming from the TBI site, it appears that the cosine similarity method is able to bypass these errors. More information on the origin of the signal could be obtained following compound annotation.

**Figure 10.**
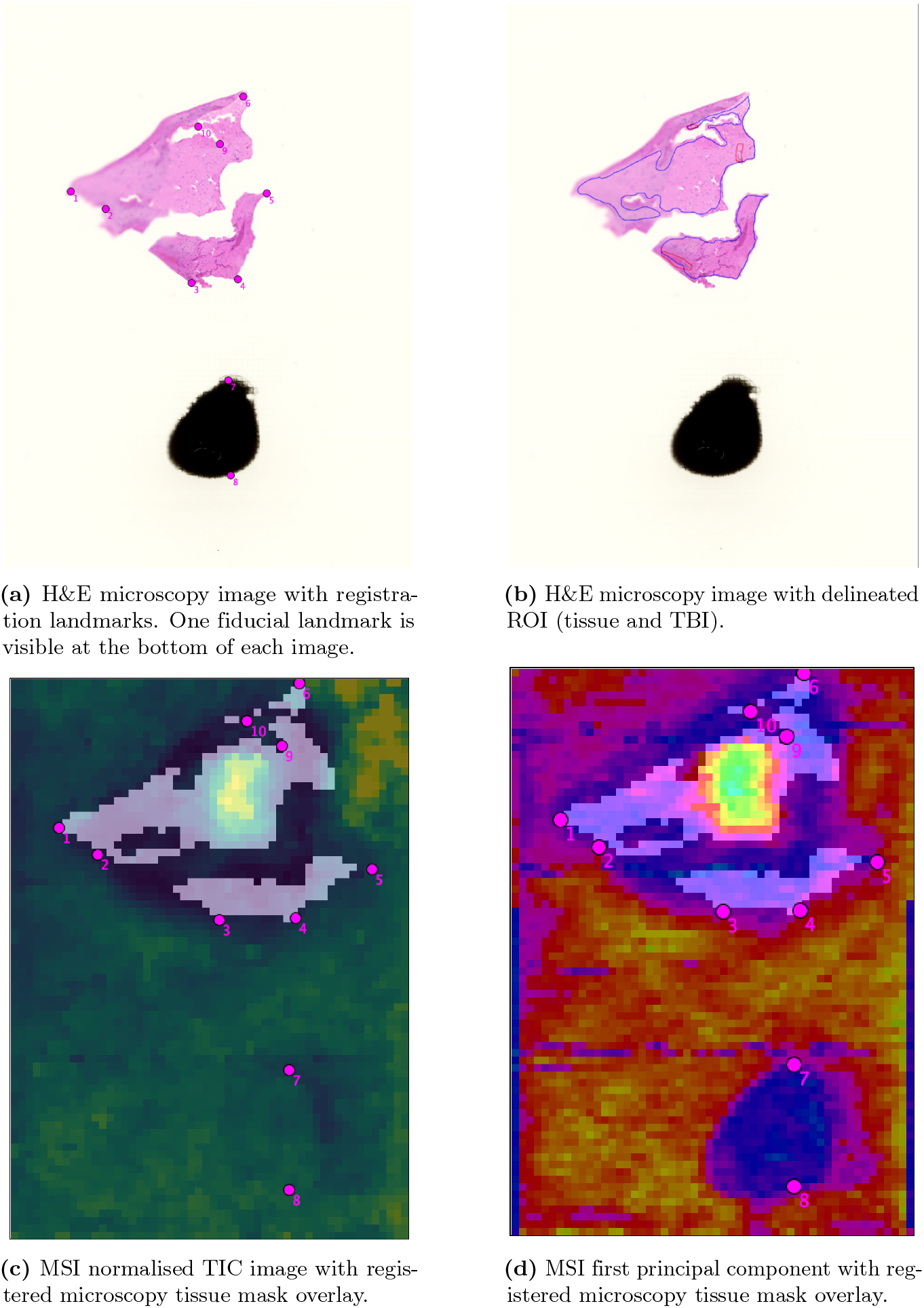
Registration results for sample 8

Sample 13 (Figure 7b) contains substantially more filtered ions, including some that appear downregulated in the TBI region. In this instance, applying a threshold based on diff_*i*_ could further refine the dataset to identify relevant ions. While the standard deviation remains relatively low for these samples, Sample 20 (Figure 7c) exhibits considerably higher standard deviation. This elevated variability may indicate either tissue heterogeneity or a more extensive TBI effect on the control region than anticipated.

## Conclusion

A method for identifying ions related to histologically annotated TBI regions in clinical MALDI-MSI datasets of brain tissue biopsies was developed. This is a proof-of-concept workflow which used a combination of methods was used ranging from registration to image co-localisation. In this case, landmark based registration was preferred due to the nature of the samples. Otherwise, automatic registration could be performed by using, for example, the MSIr package [23] by using the MSI image as the fixed image and the H&E as the moving image as opposed to their default setup.

Generally image co-localisation is used for searching ions with a similar intensity distribution pattern using an ion image as a reference. In this case, however, a binary mask was used as a reference for identifying ions with a similar intensity distribution pattern as the binary mask. Using binarised mass channels has been previously used for MSI data mining [18]. This method (Figure 1B) was validated using a synthetically generated MSI dataset. This was created by using the binary masks from the TBI dataset to assign varying intensities to the different regions subsequently followed by the addition of Poisson noise to the whole ion image [38].

Based on the results obtained from the synthetic dataset, the cosine similarity score performs better for upregulated ROI, which is to be expected. Unless the downregulated ROI is much less intense than the control region then, the similarity score will be quite similar to that of the control mass channels. Another method was to determine the cosine similarity score using the inverse of the scaled ion image for downregulated ROI mass channels. However, this method did not provide better results for detecting ions with downregulated ROIs.

Due to the fact that cosine similarity might be influenced by pixels with higher intensities, it might miss ions with lower pixel intensities in the ROI region. At low signal to noise ratios, both the magnitude and the direction of the vector become more susceptible to noise. So, the difference between the intensities in the different brain regions was also taken into consideration. Bootstrap sampling was applied to sample the control region in the brain in order to get a better estimate of the mean intensity across the tissue which excludes the TBI region. This method accounts for both the difference in size between control and TBI regions, where a specific control area is not delineated in the tissue, and tissue MSI heterogeneity. Further validation of this methods could be achieved by selecting a ROI for which the MS fingerprint is known and using that as a reference dataset.

The synthetic dataset generation could also benefit from further optimisation. For example, this can be improved by using generative adversarial networks (GANs) which have been applied to the generation of synthetic medical imaging data [17]. Once this is achieved, the synthetic datasets can be created for each dataset using the binary masks and can be further used as training datasets for machine learning (ML) models.

Further possible steps that could be undertaken for a similar study are listed next. In this case, the lists of the filtered ions for all samples were checked for any intersection. However, for samples with better signal, it would be recommended to first align the MSI samples and then determine characteristic MS features for TBI biology based on the different groups they are classified into by training a ML algorithm. Additionally, combining the results from the similarity and the difference could also be possible by Gaussian weighting the results for each method and combining the two weights for each mass channel. Finally, annotation could be easily performed by mapping the aligned dataset to an annotated tandem liquid mass spectrometry (LC-MS/MS) dataset obtained on the same MS platform from the pooled tissue homogenate samples. The same platform should be used for an increased annotation accuracy, as it helps maintain a similar mass resolution.

In conclusion, a simple but novel workflow was developed for identifying *m/z* features associated to TBI in clinical MALDI-MSI datasets. The workflow’s applicability could extend to clinical studies with aims similar to the presented in this paper.

## Data Availability

The python scripts used for this study are available on GitHub: https://github.com/mdcatapult/marshmallow-msi.

## Authors Contributions

- Conceptualization: A-M.N.
- Methodology: A-M.N. (algorithm development, computational pipeline), I.O. (MSI sample acquisition, tissue sample preparation and histology), P.K.Y. (histological annotation).
- Resources: P.K.Y., C.E.G.U. (clinical samples collection and securing funding)
- Writing – Original Draft: A-M.N.
- Writing – Review & Editing: A-M.N., P.K.Y., C.E.G.U., I.O., H.B.
- Supervision: H.B.

## Acknowledgments

The authors would like to thank Barts Charity for funding the collection of the clinical samples [45] and Joshua Millar for his contribution to the MSI method development and experimental work.

## 1 Supporting Information

### 1.1 Sample preparation and MALDI-TOF imaging

In total, 24 samples were analysed and processed using the workflow presented in this paper. The biopsy samples were provided by Queen Mary University of London (QMUL) and Barts Health NHS Trust and they come from patients who suffered a TBI of different severity forms. Based on the GOS-E scale [49], the samples could be categorised in 3 different groups. The dataset itself is confidential and only certain technical aspects regarding it can be disclosed. The tissues were formalin-fixed sucrose-embedded (FFSE). Tissue size on average 1.5 by 1.5 *µm*.

The MALDI-TOF MSI dataset was acquired using a Bruker RapiFlex Tissue Typer mass spectrometer in positive ionisation mode. The matrix used was *α*-Cyano-4-Hydroxycinnamic acid (CHCA). The images were generated at a resolution of 30.3 *µm*.

### 1.2 Sample preparation

Sections from clinical TBI biopsies were produced with a cryostat at 10 µm of thickness. They were thaw-mounted onto conductive Indium-Tin-Oxide coated glass slides (QMUL). Frozen sections were kept in the −80°C until analysis.

#### 1.2.1 Enzymatic digestion matrix spraying

1. Samples were subject to a series of preparation steps such as: lipid removal, hydration and heat induced antigen retrieval (HIAR).
2. Following HIAR, samples were coated with a trypsin solution and left to digest overnight.
3. Following an overnight digestion, the sample was sprayed with *α*-Cyano-4-hydroxycinnamic acid (CHCA) matrix solution.

### 1.3 MALDI-MSI

Samples were analysed using a Bruker RapiFlex TissueTyper mass spectrometer in positive ion mode. The data acquisition parameters were optimised to maximise the signal intensity obtained for these samples with the highest possible spatial resolution. The acquisition parameters applied to all samples are shown in Table 1.

**Table 1.** Optimised parameters for MSI data acquisition using the Bruker Rapiflex platform.

| Parameter | Value | Units |
| --- | --- | --- |
| Laser power | 36 | % |
| Laser size | 26 | $\mu\text{m}$ |
| Resulting field size | 30 | $\mu\text{m}$ |
| Mass start | 700 | Da |
| Mass end | 1800 | Da |
| Sample rate | 5 | GS/s |
| Detector gain | 1.7 | au |
| Baseline offset adjustment | 0 | % |
| Shots | 2000 | au |
| Frequency | 10000 | Hz |

Additionally, fiducial markers were drawn manually with an alcohol-resistant marker. The purpose of these markers is to have a reference to locate and select the tissue area through the camera fitted inside the mass spectrometer for analysis. The visualisation of one of the TBI samples and fiducial markers is shown in Figure 8.

### 1.4 H&E staining and brightfield microscopy

Following analysis by MSI, the samples were histologically stained with Haematoxylin and Eosin (H&E). Samples used for MALDI experiments had the matrix removed by submerging the slides in 100% methanol for 2 minutes prior to H&E staining.

After H&E staining, the slides were scanned with the ZEISS Axioscan 7™ Microscope Slide scanner in brightfield mode and the magnification used was 40x. Each image was converted to TIFF format for pathological annotation with Fiji.

### 1.5 Synthetic dataset generation

Examples of ion images generated as part of the synthetic dataset are displayed in Figure 9.

### 1.6 TBI dataset

Additional images which illustrate the landmark-based registration process are presented in Figure 10. These also include the ROI annotated by the CNS neuroinjury expert with over 27 years experience (P.K.Y.).

### 1.7 Python project

This study used an internally developed Python project that aims to process mass spectrometry imaging data. The main object *imat* (which stands for image matrix) stores information about a mass spectrometry imaging dataset including its image size and mass channel intensities. This is the backbone of the rest of the modules of this Python project.

